# Inhibition of MLLT1 limits growth of MLL-AF4 leukaemias without killing healthy haematopoietic stem cells

**DOI:** 10.1101/2025.04.30.651417

**Authors:** Shivani Rajhansa, Nicholas T. Crump, Hwei Minn Khoo, Yavor Bozhilov, Paul E. Brennan, Oleg Fedorov, Cassandra Adams, Gillian Farnie, Thomas A. Milne, Adam C. Wilkinson

## Abstract

A major challenge in cancer therapeutics has been the identification of targets that are selectively toxic to cancer cells while displaying limited effects on healthy counterparts. Toxicities related to blood production from haematopoietic stem and progenitor cells (HSPCs) can be particularly problematic and result in patient morbidity and mortality. Within haematological malignancies, therapy response rates and patient survival vary widely between cancer subtypes, with leukaemias driven by the MLL-AF4 fusion protein associated with poor prognosis. MLLT1 has been recently identified as a key potential target in acute myeloid leukaemia. Here we evaluated a panel of leukaemia cell lines and healthy HSPCs for their sensitivity to the MLLT1 inhibitor SGC-iMLLT. We found that SGC-iMLLT strongly inhibited MLL-AF4-driven leukaemia growth in vitro and in vivo. By contrast, SGC-iMLLT did not alter in vitro colony forming potential of human HSPCs or affect long-term in vivo function of mouse HSPCs. These results suggest that SGC-iMLLT may have a promising therapeutic window in the treatment of MLL-AF4-driven leukaemias, and that further clinical development is warranted.

## Main Text

A major challenge in cancer therapeutics has been the identification of targets that are selectively toxic to cancer cells while displaying limited effects on healthy counterparts. Toxicities related to blood production from haematopoietic stem and progenitor cells (HSPCs) can be particularly problematic and can result in patient morbidity and mortality. MLLT1 (Myeloid/Lymphoid Or Mixed-Lineage Leukemia; Translocated To, 1; also known as ENL) has been recently identified as a key potential target in acute myeloid leukaemia (AML)^1,2^. MLLT1 along with the highly related MLLT3 (aka AF9) are both components of the super elongation complex (SEC), which plays a crucial role in transcriptional elongation^3^. Although the exact mechanisms of MLLT1/3 function have not been worked out, MLLT1 and MLLT3 have very similar structures and can act as adapter proteins for the rapid exchange of the same co-activator and co-repressor complexes^4^. In addition, MLLT1 is essential for the recruitment of SEC components to gene targets^5^. The acyl-lysine reader (YEATS) domain is essential for the function of both proteins, likely functioning in complex formation as well as chromatin anchoring^6,7^. Of the two proteins, only MLLT1 has been shown to be essential for AML growth^1,2^, but MLLT3 has recently been shown to have a key role in stimulating fetal haematopoietic stem cell (HSC) self-renewal^8^.

Because of this potential as a therapeutic target, several small molecule inhibitors (including SGC-iMLLT^9^) have been developed to inhibit the YEATS domain, targeting both MLLT1 and MLLT3^10-15^ and displaying in vivo efficacy in AML^14,15^. MLLT1-specific inhibitors have limited efficacy^16^, but recent exciting work has found that MLLT1-specific proteolysis-targeting chimera molecules are effective in vivo^5,17^ as well as displaying low toxicity^5^. Although this suggests targeting MLLT1 alone may be effective, there is always the possibility that MLLT3 could play an important role in individual patient response as well as relapse. Thus, more needs to be understood about the range of leukaemias sensitive to MLLT1/3 inhibition, and whether targeting both MLLT1 and MLLT3 could have any potential toxicity.

To date, MLLT1 and MLLT3 have primarily been investigated as a therapeutic target in AML. The t(4;11) leukaemias are poor prognosis leukaemias^3,18^ caused by chromosome translocations involving the *Mixed Lineage Leukaemia* (*MLL*, also known as *KMT2A*) and *AF4* (*AFF4*) genes, creating a novel MLL-AF4 fusion protein, primarily associated with acute lymphoblastic leukaemia (ALL)^19^. SEC activity is essential in t(4;11) leukaemias^20,21^, so we hypothesised MLLT1/3 function may be essential for this subtype of ALL as well. Interestingly, MLLT3 protein levels are relatively low in t(4;11) leukaemias^22^, suggesting that MLLT1 may be the main effector of SEC activity in this leukaemia context.

Here we evaluated the sensitivity of SGC-iMLLT on a panel of leukaemia cell lines and on healthy HSPCs. We found that SGC-iMLLT strongly inhibited t(4;11) leukaemia growth in vitro and in vivo. By contrast, SGC-iMLLT did not alter in vitro colony forming potential of human HSPCs or affect long-term in vivo function of mouse HSPCs. These results suggest that SGC-iMLLT may have a promising therapeutic window in the treatment of MLL-AF4-driven leukaemias, and that further clinical development is warranted.

To initially test the hypothesis that t(4;11) leukaemias may be sensitive to specific loss of MLLT1, we engineered the human MLL-AF4-driven SEM ALL cells to insert a FKBP12^F36V^ degron tag^23^ at the N-terminus of MLLT1 (**Figure S1A**). In these cells, MLLT1 is degraded in the presence of dTAG-13 (**Figure S1B**). The FKBP-MLLT1 SEM lines proliferated and formed colony forming units (CFUs) comparable to parental SEM cells (**Figure 1A**). However, when SEM FKBP-MLLT1 lines were grown in the presence of dTAG-13, essentially all growth potential was lost (**Figure 1A**).

**Figure 1:**
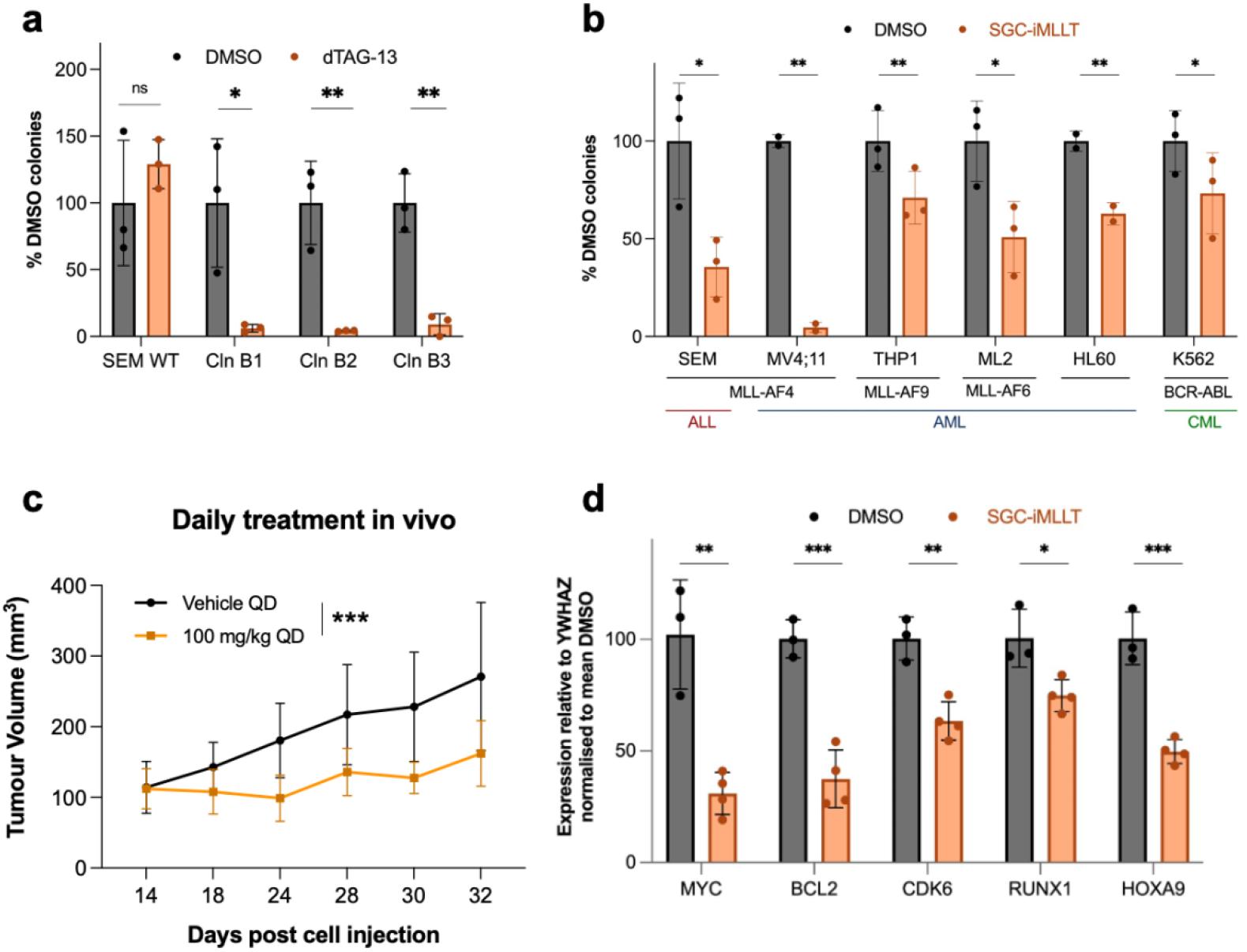
MLLT1 inhibition reduces leukaemogenic potential in vitro and in vivo. **a)** Percentage of colonies formed from SEM and FKBP-MLLT1 SEM cell lines in the presence of 1 µM dTAG-13 relative to DMSO control in three different FKBP-MLLT1 SEM cell line clones (Cln). Error bars represent s.d. of three biological replicates. Statistical analysis performed using t-tests. **b)** Percentage of colonies formed from indicated human cell lines in the presence of 1 µM SGC-iMLLT relative to DMSO control. Error bars represent s.d. of three biological replicates. Statistical analysis performed using t-tests. **c)** Tumour volume in MV4;11 xenograft model mice treated with SGC-iMLLT once a day (QD; at 100 mg/kg) for 20 days (n=9-11 mice per group). Statistical analysis performed using two-way ANOVA. **d)** RT-qPCR of MLLT1 target genes showing expression in MV4;11 xenograft-derived cells described in (c). Statistical analysis performed using t-tests. * = p < 0.05, ** = p < 0.01, *** = p < 0.001, ns = no significant difference.

Given the acute dependency of SEM for MLLT1, we next evaluated the MLLT1/3 small molecule inhibitor, SGC-iMLLT. SEM cells again lost growth potential when MLLT1/3 was inhibited (**Figure 1B**). A second human cell line carrying the MLL-AF4 translocation, AML MV4;11 cells, also demonstrated high sensitivity to SGC-iMLLT (**Figure 1B**). Other human AML cell lines, including those containing other MLL fusion proteins (MLL-AF9 (aka MLL-MLLT3), MLL-AF6), also displayed growth inhibition with SGC-iMLLT (**Figure 1B**). This indicates that MLLT1/3 inhibition may have broad activity against leukaemia.

We next asked whether SGC-iMLLT treatment could inhibit tumour growth in vivo by tracking tumour burden in immunodeficient NOD-SCID mice subcutaneously inoculated with MV4;11 cells. Daily (*Quaque die*) treatment with 100 mg/kg SGC-iMLLT significantly inhibited tumour growth over 20 days (**Figure 1C**). Additionally, analysis of the MV4;11 cells isolated at the endpoint of this assay displayed significant reductions in expression of known MLLT1 target genes (including *MYC* and *BCL2*) (**Figure 1D**). Together, these results confirm the therapeutic potential of SGC-iMLLT in t(4;11) leukaemias.

An optimal cancer therapy needs to selectively target diseased cells while showing limited toxicity for normal healthy cells. Loss of normal blood production in standard cancer treatments is particularly dangerous and is a major cause of therapy-related morbidity and mortality. We therefore investigated the effect of SGC-iMLLT on healthy human and mouse HSPCs, in order to determine the potential therapeutic window of SGC-iMLLT.

We initially assessed the in vitro colony forming potential of human CD34^+^ HSPCs, with and without SGC-iMLLT. No significant changes in colony potential were observed with SGC-iMLLT treatment (**Figure 2A, S2A**), suggesting human HSPC differentiation potential was not acutely sensitive to MLLT1/3 inhibition. Next, we assessed mouse HSCs in an ex vivo expansion culture assay. Again, no significant changes in growth were observed, and similar numbers of immunophenotypic CD201^+^CD150^+^c-Kit^+^Sca1^+^Lineage^-^ HSCs were generated in both conditions (**Figure 2B**). To understand how this compared to other leukaemia drugs, we tested the Menin inhibitor (SNDX-5613) and the CDK9 inhibitor (AZD4573) in these mouse HSC cultures. Menin inhibitor was well tolerated but we saw high toxicity from the CDK9 inhibitor (**Figure S2B**).

**Figure 2:**
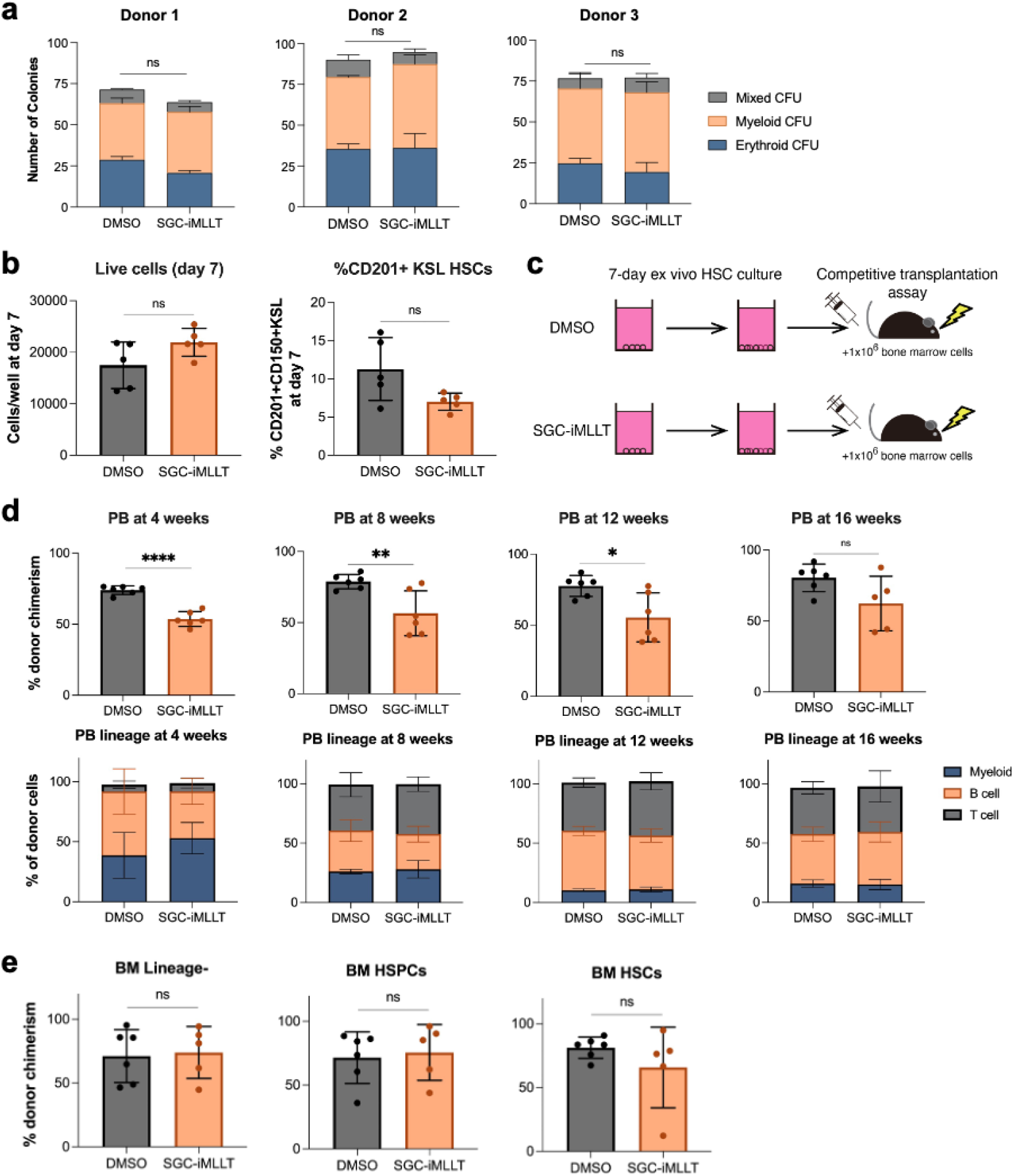
Healthy HSPCs survive MLLT1 inhibition. **a)** Number of colonies formed from 500 human CD34^+^ HSPCs grown in 1 µM SGC-iMLLT or DMSO. Erythroid, myeloid, and mixed CFUs were distinguished and counted, see **Figure S2a** for representative images. Experiments performed using 3 different donors with technical triplicates. **b)** Live cell numbers and frequency of CD201^+^CD150^+^Kit^+^Sca1^+^Lineage^-^ immunophenotypic mouse HSCs derived from day-7 HSC cultures (n=5) in the presence of 1 µM SGC-iMLLT or DMSO. **c)** Schematic of mouse HSC culture and transplantation assay to assess the effect of SGC-iMLLT on HSC activity. **d)** 4-16-week peripheral blood chimerism from 100 HSCs cultured for 7-days with 1 µM SGC-iMLLT or DMSO, transplanted alongside 1 × 10^6^ whole bone marrow competitors into lethally-irradiated recipients. Error bars represent s.d. from 5-6 recipient mice. **e)** 16-week bone marrow chimerism from 100 HSCs cultured for 7-days with 1 µM SGC-iMLLT or DMSO, and then transplanted alongside 1 × 10^6^ whole bone marrow competitor cells into lethally-irradiated recipients. Error bars represent s.d. from 5-6 recipient mice. All statistical analyses performed using t-tests: * = p < 0.05, ** = p < 0.01, **** = p < 0.0001, ns = no significant difference.

Finally, to confirm that HSC function was unaffected, we performed competition transplantation assays with HSCs treated with SGC-iMLLT for 7 days ex vivo (**Figure 2C**). While reduced peripheral blood chimerism was observed at early time points, no significant difference was observed at 16-weeks post-transplantation (**Figure 2D**). Similar lineage output was also observed in both groups (**Figure 2D**). Additionally, similar high-level donor chimerism was observed in bone marrow HSPC subsets at the 16-week endpoint (**Figure 2E**). Together, these results confirm that SGC-iMLLT has minimal toxicity on HSC activity and suggest SGC-iMLLT may have a good therapeutic window in the treatment of leukaemic blood cells, particularly MLL-AF4-driven leukaemias.

Recent studies have focused on the development of MLLT1-specific targeting molecules^5,16,17^, as MLLT1 appears to be the main effector of leukaemic growth^1,2^. However, even with highly effective targeted therapies such as inhibitors of the Menin protein, there are often patients that do not respond to treatment^24^. Expression of MLLT3 could provide an escape mechanism by which leukaemia cells can bypass the requirement for MLLT1. Thus, successful treatment may require drugs that are able to target both MLLT1 and MLLT3, which brings with it the possibility of increased toxicity. In this study, we have demonstrated that MLLT1 is required for human t(4;11) leukaemia cell lines to grow in vitro and that the SGC-iMLLT inhibitor can also limit growth in vivo. We further confirmed that SGC-iMLLT displays limited toxicity to healthy human and mouse HSPCs. This suggests that MLLT1/3 dual inhibitors may have potential as selective inhibitors of t(4;11) leukaemias, as well as having a potential wider role in some non-MLL-rearranged leukaemias.

## Figures and Legends

**Supplementary Figure 1:**
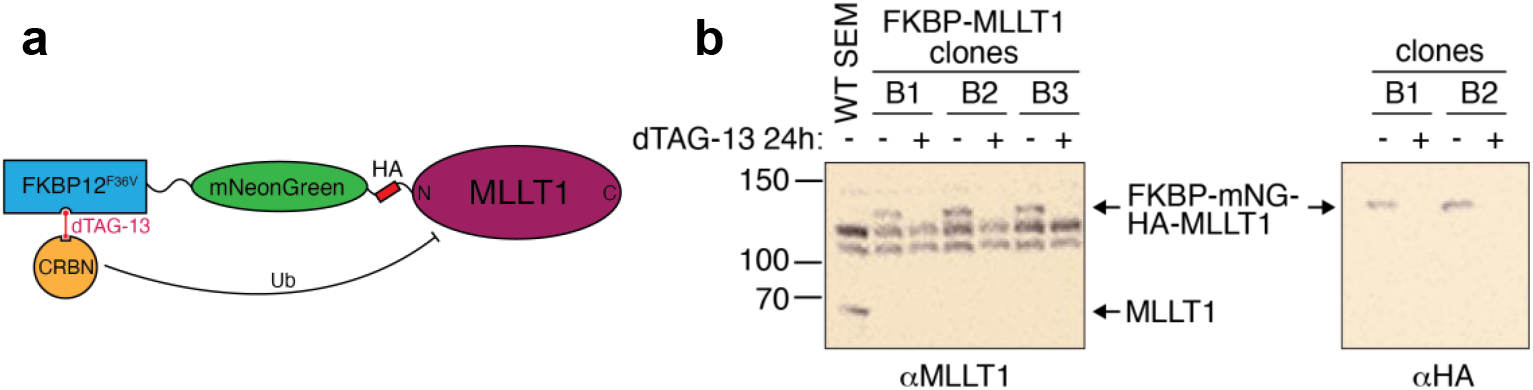
Degron tagging of MLLT1 in SEM cells. **a)** Schematic of gene editing strategy to generate MLLT1-degron tagged SEM cell lines. **b)** Western blot of wild-type MLLT1 and FKBP-mNG-HA-MLLT1 in MLLT1-degron tagged SEM cell lines used in **Figure 1a** incubated with 1 µM dTAG-13 or DMSO.

**Supplementary Figure 2:**
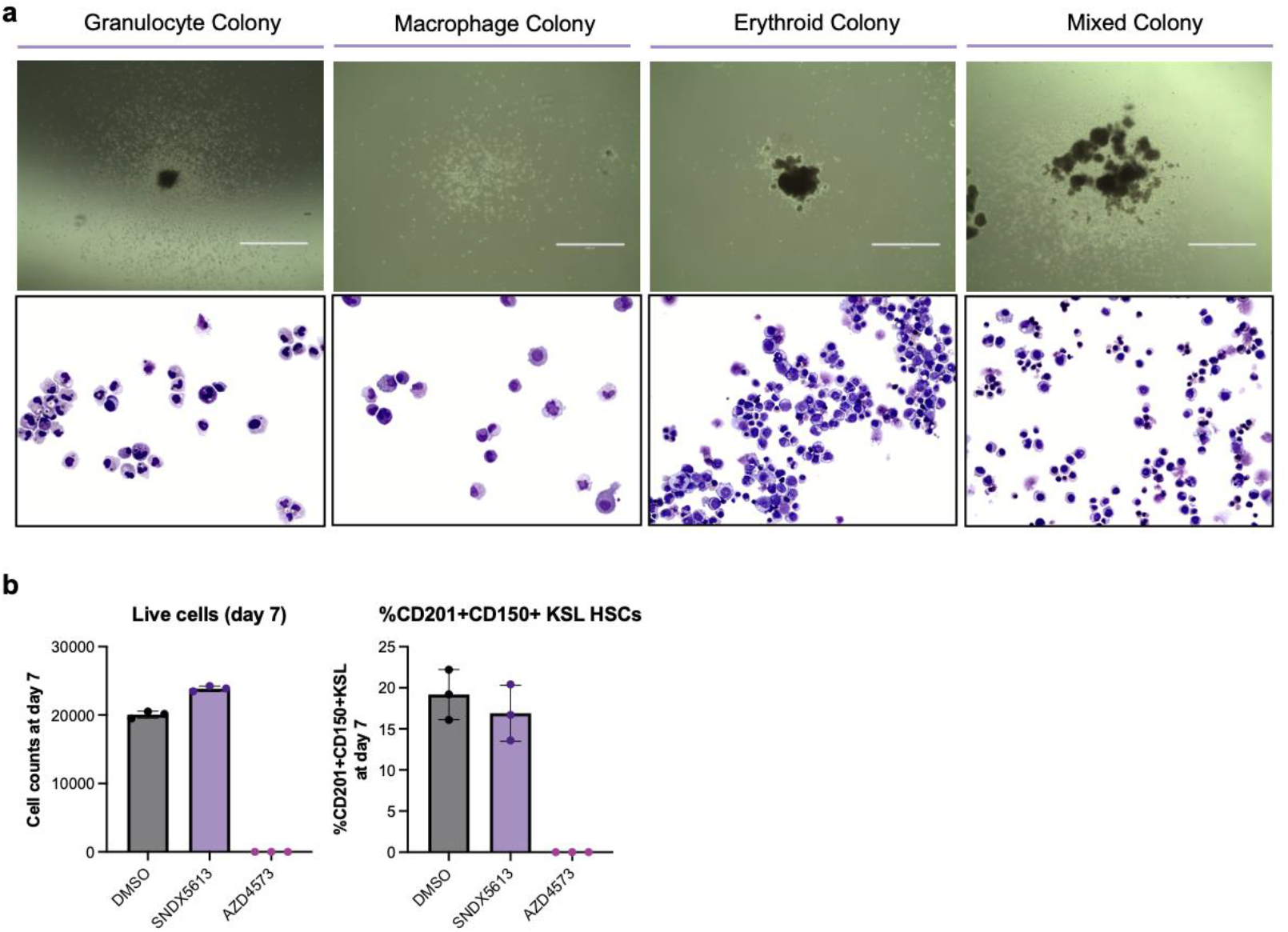
Haematopoietic stem and progenitor cell toxicity assays. **a)** Representative imaging of colonies counted in **Figure 2a** categorised as originating from a granulocyte, macrophage, erythroid, or multi-potential (“Mixed CFU”). Granulocyte and macrophage colonies counted together as “Myeloid CFU”. Upper panel shows representative imaging of colonies. Lower panel shows morphology of cells from the respective colonies. **b)** Live cell numbers and frequency of CD201^+^CD150^+^Kit^+^Sca1^+^Lineage^-^ immunophenotypic mouse HSCs derived from day-7 HSC cultures (n=3) in the presence of 250 nM SNDX5613 (Menin inhibitor), 250 nM AZD4573 (CDK9 inhibitor), or DMSO control.

## Materials and Methods

### Generation of FKBP-MLLT1 SEM cell lines

SEM cell lines were engineered to introduce *FKBP12*^*F36V*^*-P2A-mNeonGreen* immediately downstream of the start codon of *MLLT1*, using CRISPR-Cas9-mediated homology-directed repair as previously described^25^. Cells were co-electroporated with two plasmids: pX458, encoding Cas9, an sgRNA sequence targeting the N-terminal sequence of MLLT1 (F: caccGGCGCCAGCCATGGACAATC, R: aaacGATTGTCCATGGCTGGCGCC), and mRuby; and a second plasmid including the *FKBP12*^*F36V*^*-P2A-mNeonGreen* sequence, flanked by 500 bp of sequence homologous to either side of the *MLLT1* start codon. mRuby-positive cells were isolated by fluorescence-activated cell sorting after 24h, then 1-2 weeks later mNeonGreen-positive cells were isolated. Clonal populations were generated by plating cells in Methocult media, and homozygously edited clones were confirmed by PCR screening and western blotting.

### Colony Forming Assay for MLLT1 dTAG line

Wild-type SEM cells and three clones of FKBP-MLLT1 SEM cells were pre-treated with either 1 µM dTAG-13 or DMSO for 24 hours prior to plating the colony forming unit assay. 500 dTAG-13-treated or DMSO-treated cells were plated in IMDM MethoCult media (H4100; STEMCELL Technologies) with 20% fetal calf serum (FCS) in triplicates, with 1 µM dTAG-13 or DMSO added to Methocult media. Colony assay plates were incubated at 37°C and 5% CO_2_ and were counted after 13 days.

### Colony Forming Assay for leukaemic cell lines

Human leukaemia cell lines were pre-treated with either 1 µM SGC-iMLLT or DMSO for 2 hours prior to plating the colony assay. 500 SGC-iMLLT-treated or DMSO-treated cells were plated in IMDM or RPMI (depending on cell line) MethoCult media (H4100; STEMCELL Technologies) with 20% FCS in triplicates, with 1 µM SGC-iMLLT or DMSO added to Methocult media. Colony assay plates were incubated at 37°C and 5% CO_2_ and were counted after 10-14 days.

### MV4;11 xenotransplantation assay

MV4;11 xenotransplantation assays were performed by Evotec, following review by the Evotec France Ethical Committee (CE-029) and SBEA (internal Animal Welfare Body), and authorized by the French Ministry of Education. 5×10^6^ MV4;11 cells were subcutaneously injected into immunodeficient SCID mice. SGC-iMLLT1 was administrated by ip route at 50 mg/kg B.I.D or at 100 mg/kg QD dosing *vs* the vehicle treated mice, with daily treatment of the mice starting when the tumour reached a mean volume of 100-150 mm^3^, after a randomization of the mice on this read out in order to constitute the groups. Tumour volume was measured very 2-3 days using digital calipers assuming an ovoid form of the tumour. At the endpoint, RNA was extracted from tumour tissue using a QIAshredder column and the RNeasy kit (Qiagen), with on-column DNase I digestion. RNA was reverse-transcribed using Superscript III (ThermoFisher Scientific) and cDNA was quantified by Taqman qPCR (ThermoFisher Scientific), using *YWHAZ* for gene expression normalization. Taqman probes: MYC, Hs0015348_m1; *BCL2*, Hs00608023_m1; *CDK6*, Hs01026371_m1; *RUNX1*, Hs00231079_m1; *HOXA9*, F: AAAACAATGCCGAGAATGAGAGCG, R: TGGTGTTTTGTATAGGGGGACC, Taqman probe: CCCCATCGATCCCAATAACCCAGC; *YWHAZ*, Hs03044281_g1.

### Human HSPC colony forming unit assays

Human primary cell experiments were approved by the University of Oxford’s research ethics committee (OxTREC-574-23). 500 CD34^+^ umbilical cord blood-derived HSPCs (purchased from NHSBT) were plated in MethoCult media (H4230; STEMCELL Technologies) in triplicates with 1 µM SGC-iMLLT or DMSO. Colony assay plates were incubated at 37°C and 5% CO_2_ and were counted after 13 days. Representative colony types are displayed in **Figure S2a**.

### Mouse HSC culture and transplantation assay

Mouse experiments were approved by the UK Home Office and the University of Oxford’s local ethics committee. C57BL/6-CD45.2 were purchased from Envigo, while C57BL/6-CD45.1 and C57BL/6-CD45.1/CD45.2 mice were bred in-house. Lineage^-^ Sca1^+^c-Kit^+^CD201^+^CD150^+^ HSCs were FACS purified from bone marrow cells of C57BL/6-CD45.1 mice and cultured as described previously^26^. HSCs were cultured in F12 media containing 100 ng/ml TPO and 10 ng/ml SCF, as described previously^26^. HSC cultures were supplemented with either 1 µM SGC-iMLLT or DMSO for 7 days. Alternatively, HSC cultures were supplemented with 250 nM SNDX5613 (Menin inhibitor), 250 nM AZD4573 (CDK9 inhibitor), or DMSO control. For transplantation assays, 7-day cultured cells (derived from 100 HSCs) were transplanted alongside 1 × 10^6^ bone-marrow competitor cells from C57BL/6-CD45.1/CD45.2 mice into lethally irradiated (10 Gy) C57BL/6-CD45.2 mice. Peripheral blood chimerism analysis was performed every 4 weeks as described previously. At the 16-week endpoint, donor chimerism was measured in both peripheral blood and bone marrow by flow cytometry, as described previously^26^.

## Acknowledgements

We thank the WIMM Flow Cytometry Core for flow cytometry access. This research was supported by grants from the Kay Kendall Leukaemia Fund (KKL1443), the MRC (MC_UU_00016/6 and MC_UU_00029/6), NIHR, NIHR Oxford Biomedical Research Centre, the Wellcome Trust, and the Krishnan-Ang Foundation.

## Author contributions

NTC, ACW, and TAM conceived the experimental design. SR, NTC, HMK, YB, OF and GF carried out experiments. SR, NTC, ACW and TAM. analysed and curated the data. SR, NTC, ACW and TAM interpreted the data and wrote the manuscript. NTC, PEB, OF, CA, GF, ACW and TAM provided funding and supervision. All authors reviewed the manuscript.

